# A single cell brain atlas in human Alzheimer’s disease

**DOI:** 10.1101/628347

**Authors:** Alexandra Grubman, Gabriel Chew, John F. Ouyang, Guizhi Sun, Xin Yi Choo, Catriona McLean, Rebecca Simmons, Sam Buckberry, Dulce Vargas Landin, Jahnvi Pflueger, Ryan Lister, Owen J. L. Rackham, Enrico Petretto, Jose M. Polo

**Author notes:** These authors contributed equally to this work (Alexandra Grubman, Gabriel Chew, John F. Ouyang). Correspondence should be addressed to J.M.P., E.P. and O.J.L.R.

## Abstract

Alzheimer’s disease (AD) is a heterogeneous disease that is largely dependent on the complex cellular microenvironment in the brain. This complexity impedes our understanding of how individual cell types contribute to disease progression and outcome. To characterize the molecular and functional cell diversity in the human AD brain we utilized single nuclei RNA-seq in AD and control patient brains in order to map the landscape of cellular heterogeneity in AD. We detail gene expression changes at the level of cells and cell subclusters, highlighting specific cellular contributions to global gene expression patterns between control and Alzheimer’s patient brains. We observed distinct cellular regulation of *APOE* which was repressed in oligodendrocyte progenitor cells (OPCs) and astrocyte AD subclusters, and highly enriched in a microglial AD subcluster. In addition, oligodendrocyte and microglia AD subclusters show discordant expression of *APOE.* Integration of transcription factor regulatory modules with downstream GWAS gene targets revealed subcluster-specific control of AD cell fate transitions. For example, this analysis uncovered that astrocyte diversity in AD was under the control of transcription factor EB (TFEB), a master regulator of lysosomal function and which initiated a regulatory cascade containing multiple AD GWAS genes. These results establish functional links between specific cellular sub-populations in AD, and provide new insights into the coordinated control of AD GWAS genes and their cell-type specific contribution to disease susceptibility. Finally, we created an interactive reference web resource which will facilitate brain and AD researchers to explore the molecular architecture of subtype and AD-specific cell identity, molecular and functional diversity at the single cell level.

**Highlights:**

- We generated the first human single cell transcriptome in AD patient brains
- Our study unveiled 9 clusters of cell-type specific and common gene expression patterns between control and AD brains, including clusters of genes that present properties of different cell types (i.e. astrocytes and oligodendrocytes)
- Our analyses also uncovered functionally specialized sub-cellular clusters: 5 microglial clusters, 8 astrocyte clusters, 6 neuronal clusters, 6 oligodendrocyte clusters, 4 OPC and 2 endothelial clusters, each enriched for specific ontological gene categories
- Our analyses found manifold AD GWAS genes specifically associated with one cell-type, and sets of AD GWAS genes co-ordinately and differentially regulated between different brain cell-types in AD sub-cellular clusters
- We mapped the regulatory landscape driving transcriptional changes in AD brain, and identified transcription factor networks which we predict to control cell fate transitions between control and AD sub-cellular clusters
- Finally, we provide an interactive web-resource that allows the user to further visualise and interrogate our dataset.

**Data resource web interface**:http://adsn.ddnetbio.com

## Introduction

Alzheimer’s disease (AD) is the most common form of dementia in the elderly, and as there are currently no effective treatments, AD is one of the leading contributors to global disease burden. Genetic mapping approaches such as genome wide association studies (GWAS) have uncovered clusters of AD susceptibility genes involved in endocytic and microglial pathways associated with the development of late onset AD (LOAD) ^1–4^. Computational analyses of LOAD bulk brain transcriptomes ^5^ pointed to changes in underlying network structure controlled by gain of microglial gene connectivity and loss of neuronal connectivity in AD. Whilst a momentous advance in understanding the transcriptional network dynamics of Alzheimer’s disease, bulk analyses did not provide enough resolution to resolve the precise cellular changes that occur in such heterogeneous diseases. For example, the effects of microglial niche expansion ^6^ or neuronal loss ^7^ in contributing to the observed network structure changes in AD remain to be elucidated at the single cell level. Thus, there is a clear need to examine the single cell landscape of AD brains in order to understand how gene regulatory networks drive transcriptional changes underlying altered cell identity in AD. Recent advances in droplet based RNA-sequencing technologies such as DropSeq ^8^ and single-nucleus RNA-sequencing (DroNcSeq) ^9^ have revolutionised the ability to capture single nuclei at unprecedented scale and process archival brain tissue. Taken together, these advances can facilitate a comprehensive analysis of cell heterogeneity and cell-specific changes occurring in AD brains. Several studies have examined the single cell transcriptional landscape during human brain development ^10, 11^, although to date no study has examined global cell diversity in a human brain disease.

Here, we applied an unbiased approach using DroNcSeq to characterise the cellular heterogeneity of AD patient brains in the affected entorhinal cortex. We identified novel sub-populations of cells which were present only in AD brains and with common and distinct networks of coregulated genes and functions across different cell types. Importantly, as a proof of principle for the utility of this data resource, we mapped GWAS genetic data onto these regulatory modules and identified how GWAS genes may functionally influence AD susceptibility in specific cell subclusters and to aid discovery of possible pathogenic subclusters and cell fate transitions underlying AD. Lastly, we created a searchable web interface for this AD brain cell atlas, providing a useful resource to dissect mechanisms of cell heterogeneity and dysfunction in AD.

## Results

### A molecular survey of the human AD brain

To investigate cell diversity and disease-related cellular changes in network structure in AD, we performed DroNcSeq on the 10X platform (Fig. 1a, Extended data Fig. 1). After quality control filtering, we obtained 13,214 cells with median 646 detected genes per cell from *n*=6 AD patients and *n=*6 sex and age-matched controls (mean age 77.6, range 67.3-91 years, Extended data Fig. 2, Supplementary Table 1), including a range of *APOE* genotypes (E3/3, E3/4, E4/4, E2/4). Visualization of single nuclei transcriptomes in UMAP space was able to separate nuclei into clusters which we mapped to the 6 *a priori* cell types (microglia, astrocytes, neurons, oligodendrocyte progenitor cells, oligodendrocytes and endothelial cells) based on previously established cell type specific gene sets (^12^, Fig. 1c). We defined a cell type score for each gene set and classified cells as “hybrid” if the two highest cell type scores were within 20% of each other, whereas cells within the lowest 5% of cell type scores for each cell type were defined as “unidentified” (see Methods for details). Although there is a possibility that after filtering and QC (in which doublets were excluded), hybrid cells may still represent doublets, these cells may represent true intermediate cellular states ^13^, and hence we retained them for investigation. We detected a high proportion of oligodendrocytes compared to neurons and astrocytes, and also compared to previous DroNcSeq studies ^9, 14^. Consistent with our findings, a recent study of regional cell density in the human brain revealed that approximately 40% of cortical grey matter is composed of neurons, whereas neurons only comprised ∼15% of the hippocampal formation, which included entorhinal cortex, and this proportion was further reduced to approximately 5% in AD ^15^. However, this result needs to be taken in the context that the unequal cell proportions observed between studies may reflect tissue sampling differences or sample preparation techniques. For instance, where previous studies sampled either Brodmann areas 6,9,10,17 and cerebellum ^14^, or hippocampus and prefrontal cortex ^9^, our study sampled entorhinal cortex.

**Fig. 1.**
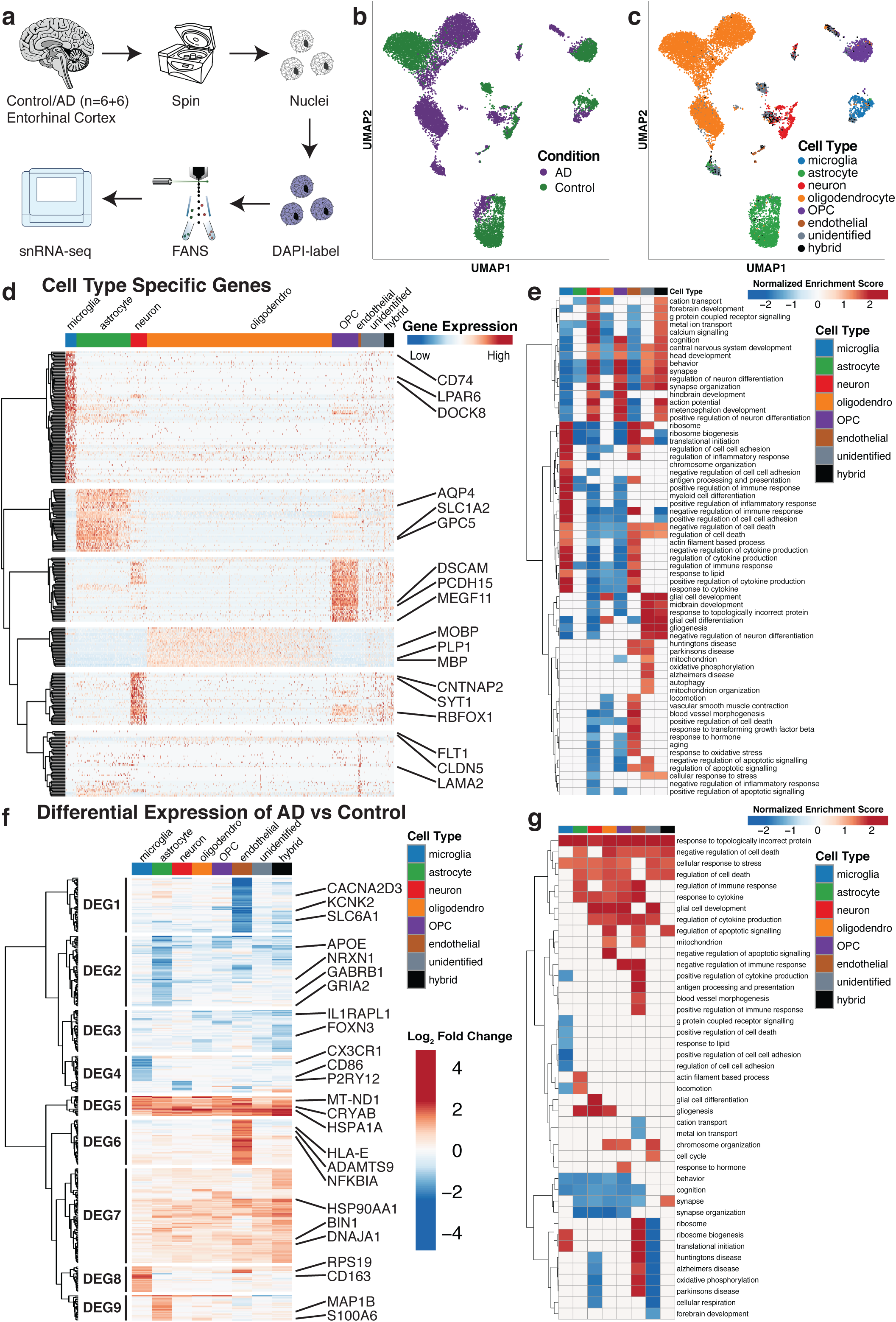
Single nuclei sequencing of human AD and control entorhinal cortex recapitulates cell types specific marker genes and cell type specific changes in AD. **a**, schematic of nuclei isolation and RNA-seq workflow. **b**, UMAP visualisation showing clustering of single nuclei, coloured by the disease diagnosis, or **c**, cell types, based on our scoring system using the gene sets determined by BRETIGEA^12^. For each cell, cell type scores were calculated separately for the six cell types in BRETIGEA and a cell is assigned the cell type with the highest score. Furthermore, we defined unidentified cells to be cells with low scores across the six cell types tested and defined hybrid cells to be cells where the top two highest cell type scores are similar. **d**, Hierarchical clustering and heat map coloured by single cell gene expression of cell-type specific genes. **e**, Gene set enrichment analysis (GSEA) of cell-type specific genes coloured by normalised enrichment scores. **f**, Hierarchical clustering and heat map showing LFC of DEGs between AD and control cells for each cell type. **g**, GSEA of the differential expression between AD and control for each cell type coloured by normalised gene set enrichment scores.

**Fig. 2.**
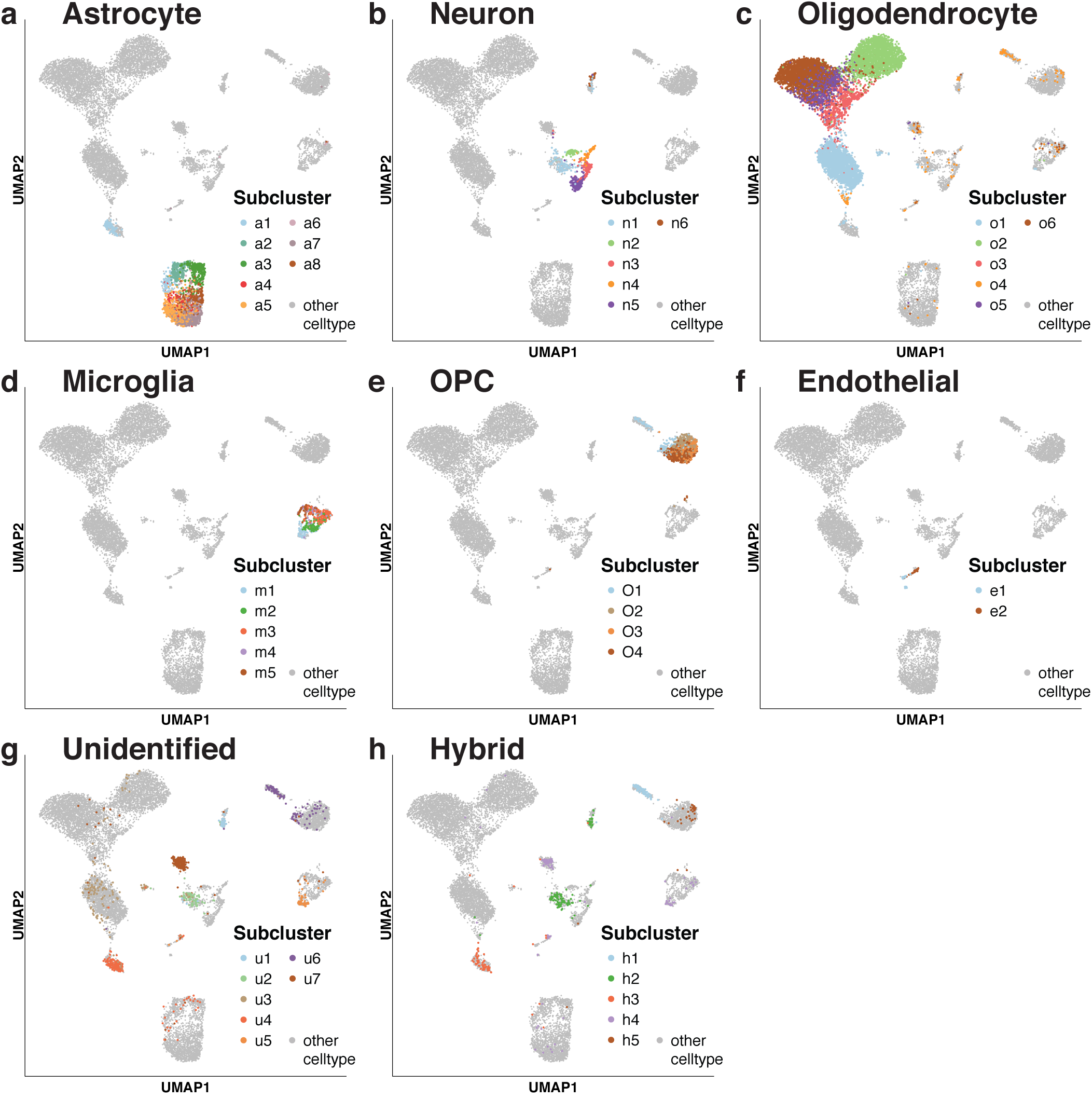
Single nuclei sequencing of human entorhinal cortex uncovers high cellular heterogeneity with each cell type. Global UMAP visualisation showing the location of cells within each cell subcluster of astrocytes (**a**), neurons (**b**), oligodendrocytes (**c**), microglia (**d**), OPCs (**e**), endothelial (**f**), unidentified (**g**) and hybrid (**h**) cells. The cell types were determined previously in Figure 1c.

In UMAP space, cells from AD patients segregated from those in controls for each cell type (Fig. 1b). We then identified marker genes for each cell type (Fig. 1d), which included *bona fide* marker genes such as *HLA-DRA, CX3CR1, C1QB*, and *CSF1R* for microglia*, AQP4, SLC1A2* for astrocytes, *SYT1*, glutamate receptors *(i.e., GRIK2, GRIA1* and *GRIN2B), RBFOX1* for neurons, *MOBP, MBP* and *PLP1* for oligodendrocytes, *PCDH15* and *MEGF11* for OPCs, and *FLT1* and *CLDN5* for endothelial cells ^10, 16, 17^. In addition, our analyses identified several putative novel specific cell-type marker genes (Supplementary Table 2), including *KCNQ3* in microglia, a Voltage-Gated Potassium Channel that controls neuronal excitability, with multiple previously described mutations causing benign neonatal epilepsy ^18^. We also identified a novel astrocyte-specific gene, *ADGRV1*, encoding the nervous system-restricted calcium binding G coupled protein receptor, GPR98, mutations in which cause Usher disease_19._

We confirmed that the functional annotation of the cell-type specific gene sets was largely consistent with the biological function of those cells (Fig. 1e). Neurons and OPCs were enriched for genes related to action potential, cognition and synaptic function, whereas microglia and endothelial cells were enriched for ribosome biogenesis, cytokine and lipid response functions, as well as antigen processing and presentation. Individually, neurons showed enrichment for genes involved in calcium signalling and metal ion transport, endothelial cells for blood vessel morphogenesis and microglia for myeloid cell differentiation and inflammatory responses.

We then set out to examine the cell type specific gene expression changes between control and AD nuclei in the context of existing literature. Our analyses unveiled nine clusters of cell-type specific and common gene expression patterns between control and AD brains (Fig. 1f, g). The cell types with the most coordinated gene expression differences between control and AD patients were microglia (further analyses reported in Grubman et al., accompanying manuscript), astrocytes and endothelial cells. These coordinated changes in expression can be summarised as clusters of down- and/or up-regulated genes (DEG) in specific cell-types, *e.g.*, clusters DEG1 and DEG6 in endothelial cells, DEG2 and DEG9 in astrocytes and DEG4 and DEG8 in microglia (Fig. 1f). We also found clusters of genes that were coordinated in their induction (DEG5 and DEG7) or repression (DEG3) in AD patient brains across multiple cell types. For instance, both DEG5 and DEG7 clusters of highly and co-ordinately upregulated genes in AD brains were enriched for genes involved in responses to topologically incorrect protein and cell stress responses, including a number of mitochondrial, heat shock and chaperone genes (*i.e., MT-ND1-4, MT-CO2, MT-CO3, MT-ATP6, HSPA1A, HSP90A1, DNAJA1* Fig. 1f, g), consistent with previous reports in SH-SY5Y cells and primary mouse neurons ^20^. It is possible that the cell-independent responses to topologically incorrect protein may be responses to extracellular amyloid deposition, as tau is only intraneuronal and there is evidence of plaque and oligomer interactions with multiple brain cell types ^21^. Our analyses highlighted that, in addition to the well-studied cell-type-specific responses (*i.e.*, inflammatory responses of astrocytes and microglia in AD, oxidative stress in neurons in AD ^22–24^), there is a further coordinated response which may act to boost molecular chaperone levels to protect cells against protein misfolding and which may provide additional therapeutic avenues aimed at enhancing endogenous cell-type independent responses.

Endothelial cells in AD upregulated genes involved in cytokine secretion and immune responses including *HLA-E, MEF2C* and *NFKBIA*. Endothelial cells and microglia in AD were enriched in ribosomal processes and translation initiation (*i.e.*, *RPS19, RPS28*). The DEG2 module of genes, which includes GABA receptors (*GABRA2, GABRB1*), glutamate receptors (*GRIA2, GRID2*) and neurexin genes (*NRXN1* and *NRXN3*), was enriched in functions related to behaviour, cognition and synapse organization functions and was downregulated in AD neurons, astrocytes, oligodendrocytes and OPCs. This is consistent with a previously reported loss of synapse module connectivity occurring in the AD prefrontal cortex ^5^. We also found AD microglia downregulated homeostatic genes (*i.e., CX3CR1, P2RY12, P2RY13* from DEG4) as has been previously described in multiple AD mouse models ^25, 26^, (Grubman et al., accompanying manuscript). Furthermore, AD microglia also downregulated genes related to cell-cell adhesion (*i.e., CD86, CD83*), lipid response (*LPAR6*) and G-coupled protein receptor pathways (*CNR1, GPR183, VIP, LPAR6*). We observed that genes related to glial cell development and differentiation and in particular the organisation and control of myelination were highly upregulated in AD neurons, astrocytes and oligodendrocytes (*i.e., BIN1* ^27^*, CNTN2* ^28^*, NDRG1* ^29^). These changes in gene expression may also reflect compensatory responses to the myelin loss observed in AD (reviewed in ^30^), although this protective mechanism in APP/PS1 mice was previously not replicated in AD post mortem tissue by staining for protective OLIG2^+^ precursors ^31^. Oligodendrocytes, astrocytes, OPCs and endothelial cells in AD were enriched for genes involved in negative regulation of cell death, suggesting a coordinated response to compensate for cell stress pathways and protect damaged cells ^20^.

Interestingly, while endothelial cells in AD upregulated genes involved in processes related to neurodegeneration (Huntington’s, Alzheimer’s and Parkinson’s diseases) and oxidative phosphorylation, we discovered that AD neurons show a downregulation of genes associated with neurodegeneration. We thus examined the cell-specific DEGs in our dataset annotated as associated with AD in the Kyoto Encyclopaedia of Genes and Genomes (KEGG) database ^32^, and discovered that the majority (*i.e.*, 60% in neurons, Supplementary Table 3) of DEGs contributing to the AD ontology enrichment are nuclear encoded mitochondrial complex I to V genes of the *NDUF, SDH, UQCR, COX*, and *ATP5* families, suggesting increased respiratory potential in these cell clusters.

Given the known specific vulnerability of cholinergic neurons in AD ^7^, we investigated whether specific regulatory pathways were different in excitatory and inhibitory neurons in AD. We mapped our single cells from each neuronal subcluster onto the excitatory (Ex1-8) and inhibitory (In1-8) neuronal signatures described in ^33^ (Extended data Fig. 3). Our results showed downregulation of specific excitatory neuron genes related to synaptic transmission (*i.e.*, *SNAP25, RIMS1*), and downregulation of ion transport and learning/memory-associated genes that was specific to inhibitory neurons (*i.e.*, *CCK, SST, RELN, VIP, KCNIP4*; Extended data Fig. 3). Together these analyses provide a detailed snapshot of both global and cell-type specific changes in gene expression and functional processes underlying AD in human brain.

**Fig. 3.**
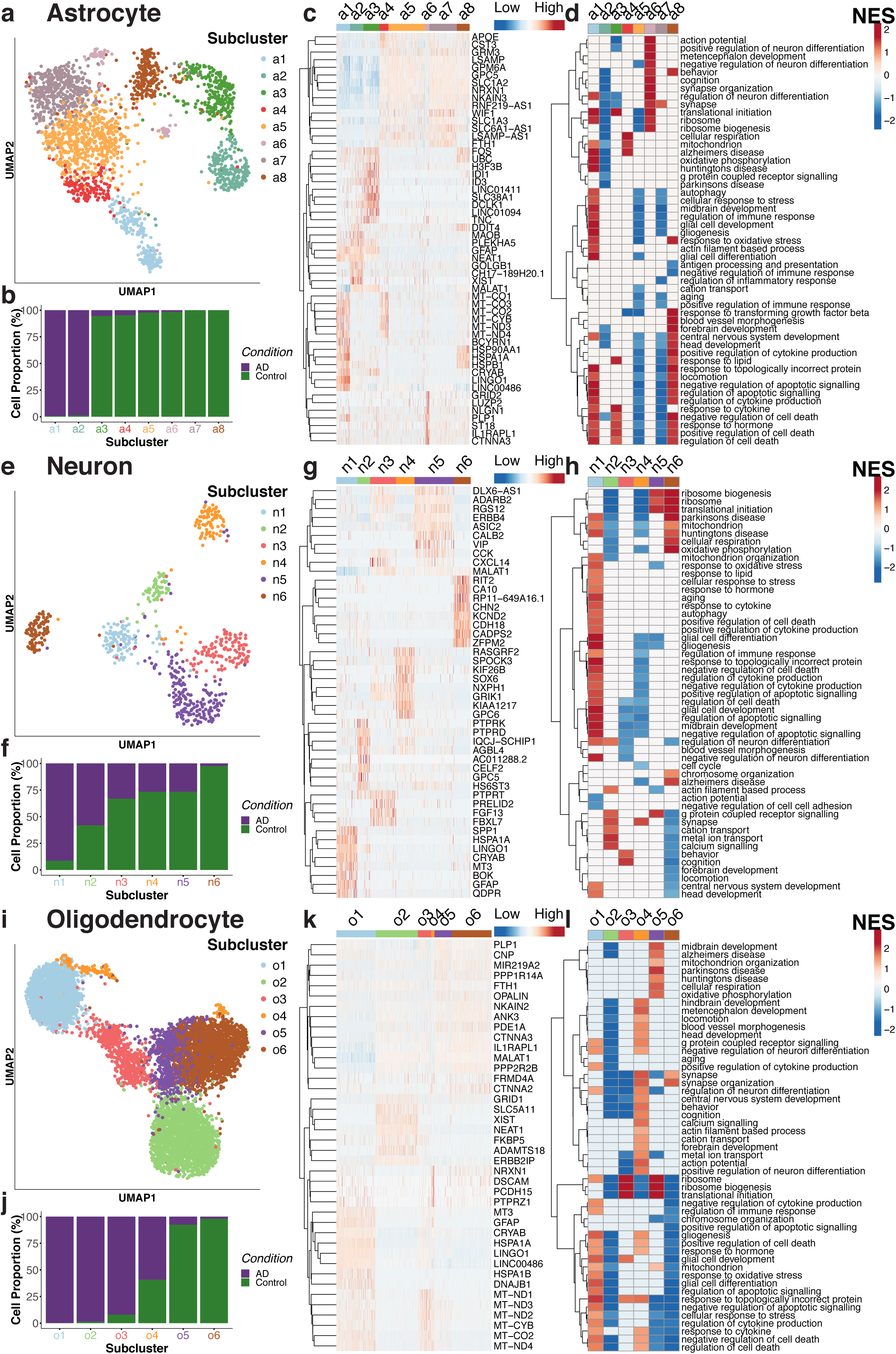
Single nuclei sequencing of human AD and control entorhinal cortex reveals homeostatic, AD-specific and shared ontological cell subclusters. a,e,i, UMAP visualization of subclusters of astrocytes (**a**), neurons (**e**) and oligodendrocytes (**i**) showing **b, f, j**, the composition of cells in subclusters by disease state, **c, g, k,** hierarchical clustering and heat map colored by single cell gene expression of subcluster-specific genes (top 8 genes were shown per cluster). **d, h, l,** GSEA of subcluster specific genes coloured by normalised gene set enrichment scores for the gene ontologies shown in each cell subcluster.

### Subcluster-specific analysis identifies homeostatic, AD-specific and shared ontological cell subclusters

As our analyses showed evidence for substantial cellular heterogeneity, we next used the Seurat algorithm ^34^ to obtain subclusters within each cell type (see Methods for details). This analysis uncovered five microglial clusters, eight astrocyte clusters, six neuronal clusters, six oligodendrocyte clusters, four OPC and two endothelial clusters (Fig. 2, Fig. 3a, e, i), each enriched for specific functional categories (Fig. 3d, h, l, Extended data Fig. 4). Except in neurons, where four clusters were composed of a mixture of AD or control cells, we found that AD and control cells mostly segregate into different clusters (Fig, 3b, f, j, Extended data Fig. 2b), suggesting strong and penetrant disease-associated transcriptional changes across almost every cell type.

**Fig. 4.**
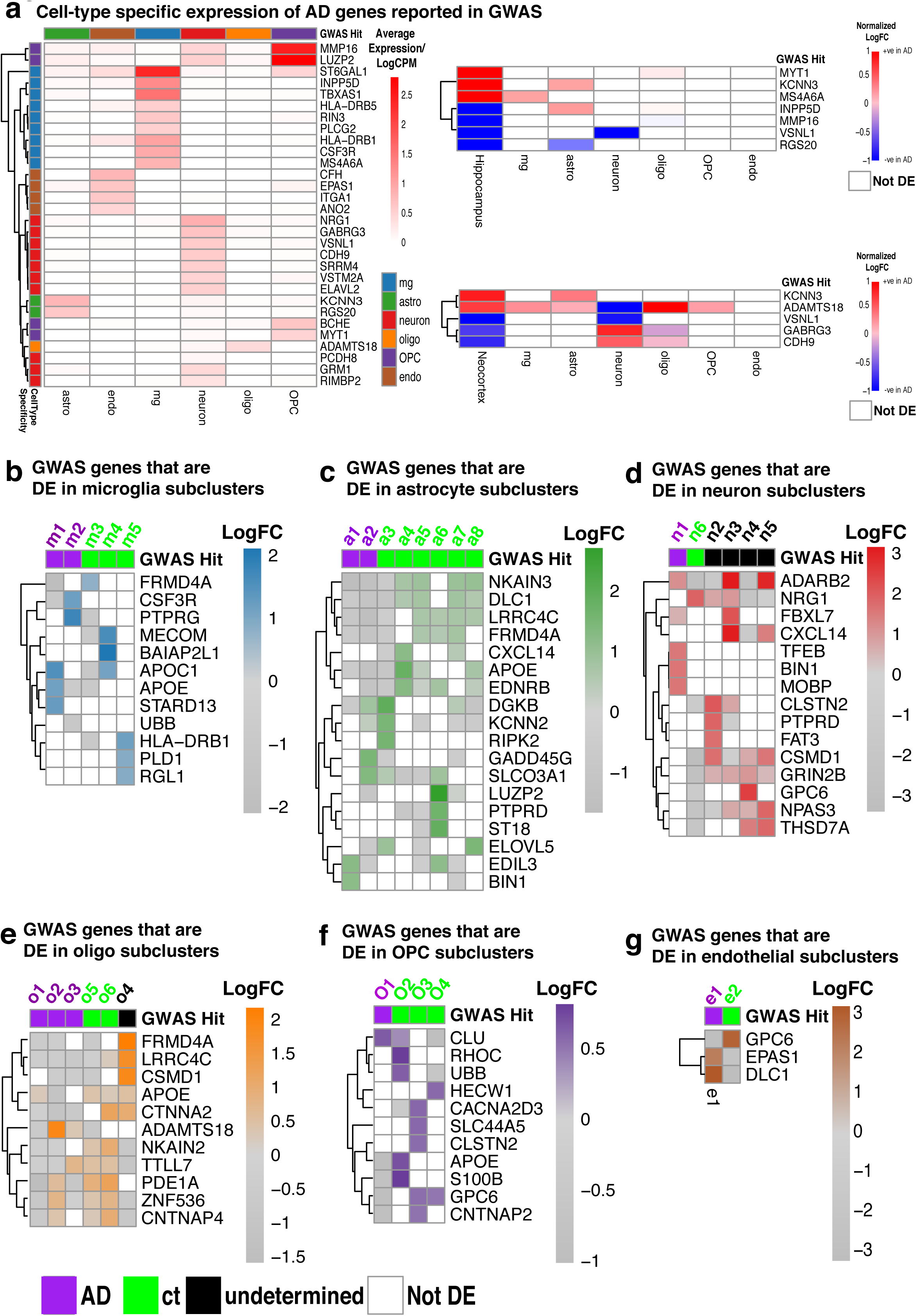
AD genes identified by GWAS show specific gene expression patterns across cell-types and cell type subclusters. a, *left panel*, Expression of cell-type specific GWAS hits (*n*=30, >99th % for specificity) and, *right panels*, comparison of cell-type specificity of GWAS DEGs identified by previous bulk tissue microarray studies between control and AD brains. **b-g,** Heatmaps showing differential expression (LFC) for GWAS genes (FDR<0.05) within subclusters with respect to each cell type; for each subcluster, the top 3 GWAS genes based on absolute log fold change were chosen for visualization; microglia (**b**), astrocytes (**c**), neurons (**d**), oligodendrocytes (**e**), OPCs (**f**) or endothelial cells (**g**). Subclusters were considered as AD or control subclusters if >80% of cells in that subcluster were from either AD or control brains respectively, otherwise they are considered undetermined.

One of the AD astrocyte subclusters, a1, was located closer to oligodendrocytes than to astrocytes on the global UMAP space (Fig. 2a). This a1 AD astrocyte subcluster was enriched for ribosomal, mitochondrial, neuron differentiation and heat shock responses, whereas the a2 AD astrocyte subcluster showed downregulation of these processes (Fig. 3d), and was instead enriched for TGFβ signalling and immune responses (refer to our web resource for full list of gene set enrichment). Neither the a1 nor the a2 AD astrocyte subcluster described here overlapped significantly with the A1 or A2 astrocyte profiles previously described in mouse AD brains, although the marker gene *C3* was upregulated in AD astrocytes as previously reported (^24^, Extended data Fig. 5a), specifically in the presently defined a2 subcluster. We noticed that in healthy patient brains, molecularly distinct astrocyte subclusters appeared to show evidence of functional diversity, as a4 was enriched in respiratory and mitochondrial processes, a3 and a8 were enriched in cellular responses to lipids and hormones, whereas a6 was enriched in synapse organisation, action potentials and ion channel activity. This suggests functional specialisation towards immune surveillance or synaptic signalling functions in individual astrocytes (^35^, Extended data Fig. 5a_i_).

**Fig. 5.**
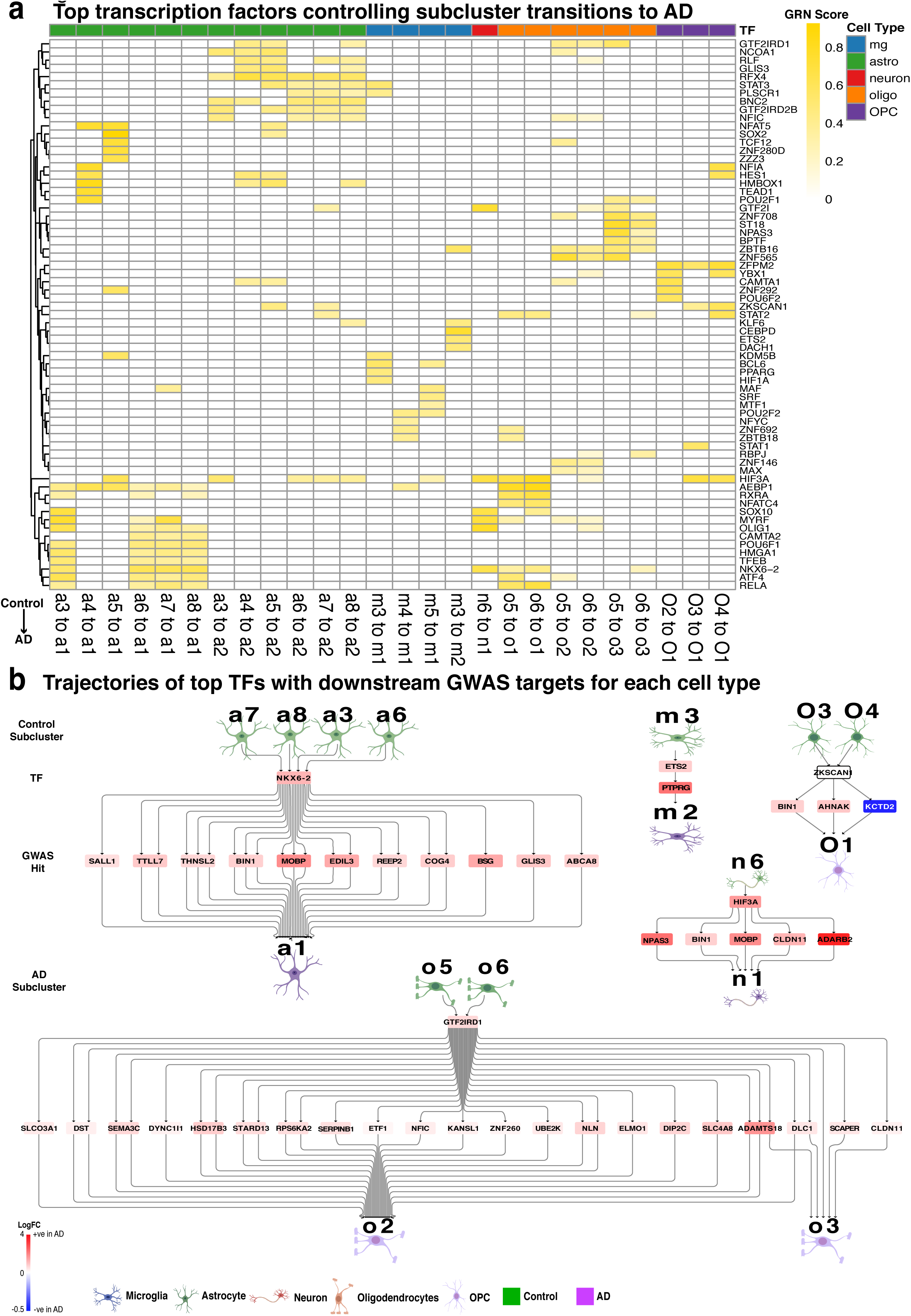
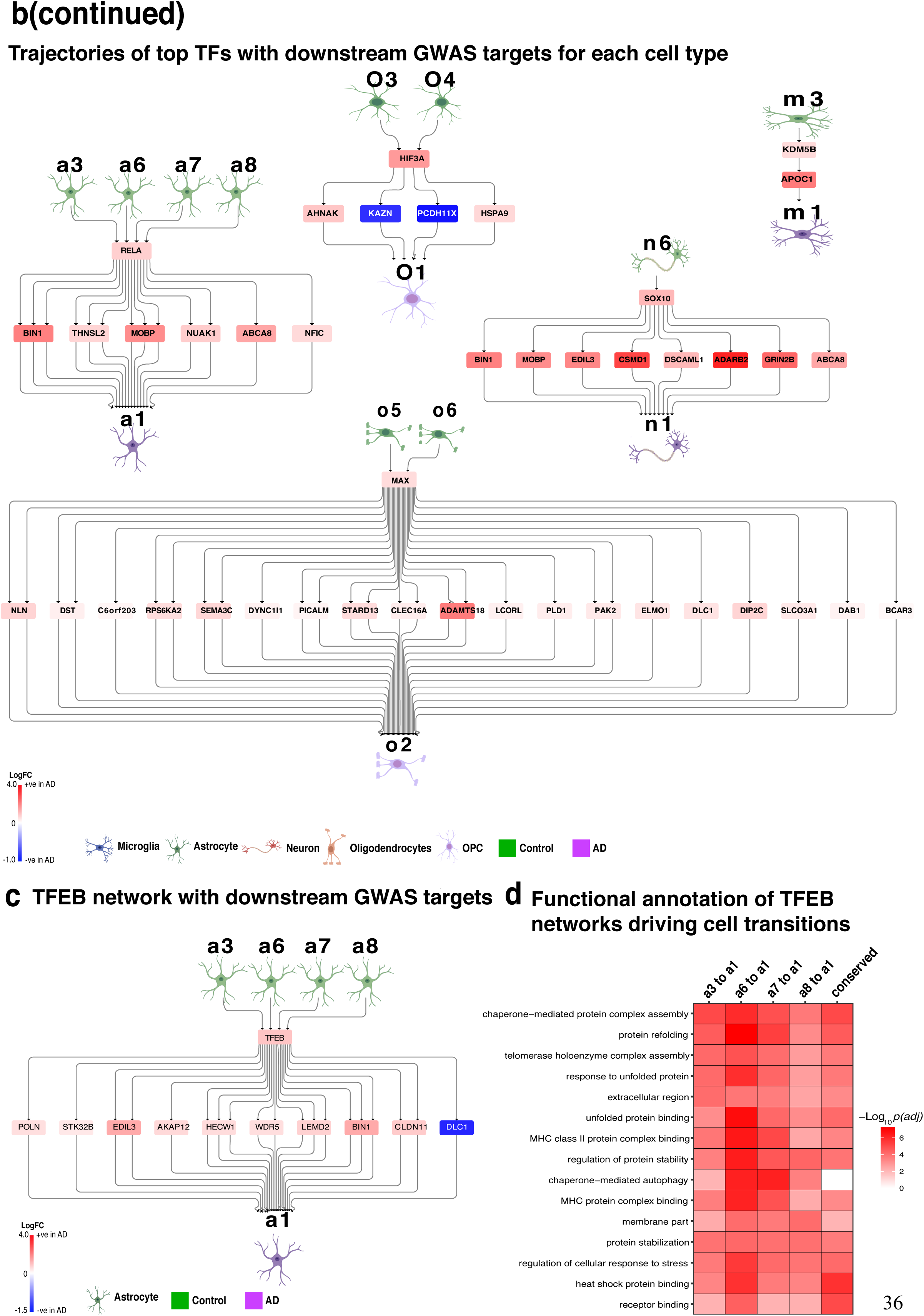
Gene regulatory network analysis predicts transcription factors for conversion of control to AD subcluster signatures. **a**, Heatmap of scores from top regulons for cell fate transitions from control subclusters to AD subclusters. Note that some transitions such as m4 to m2, m5 to m2 are not shown, as no significant TF regulons were identified after post-processing of CellRouter outputs (See Methods). **b**, Trajectories of top TFs with downstream GWAS targets for each cell type. **c,** TFEB network with downstream GWAS targets. **d,** Functional annotation of TFEB-networks driving cell transitions

We next investigated subcluster-specific changes occurring in AD, and to this aim we performed differential expression analysis between a subcluster of interest and the remaining subclusters of the same cell type. We observed some instances of genes implicated in both neurological and psychiatric disorders showing subcluster-specific differential expression in AD. For example, *LINGO1*, encoding a signalling protein that inhibits myelination ^36^, was strongly upregulated specifically in several AD subclusters (a1-a2, m1, n1, o1 and o3, Fig. 3c, g, k). Notably, *LINGO1* interacts with APP ^37^ and is a promising target for multiple sclerosis (MS) with a LINGO1 neutralising antibody in Phase II trials ^38^. Similarly, *NEAT1*, a regulatory non-coding RNA that is found increased in serum of patients with relapsing-remitting MS ^39^, was highly expressed in a1, a2 and o2 AD subclusters (Fig. 3c, k). Conversely, the glutamate transporter, encoded by *GRID1*, which has been linked to schizophrenia ^40^, was specifically increased in o2 but downregulated in the a1 subcluster (Fig. 3k). Therefore, the specific differential expression patterns shown here for the above examples can inform targeted functional studies to regulate these genes in specific subcellular compartments in AD brain.

Beyond subcluster-specific gene expression changes in AD, our data also provide insights into the molecular nature of cellular heterogeneity in AD patient brains. With respect to neuronal subclusters mapped onto excitatory and inhibitory neurons (Extended data Fig. 3), we found that clusters n2-n5 contained cells from both control and AD patient brains that separated according to functional excitatory/inhibitory identity. Cluster n2 comprised excitatory neurons corresponding to a mixture of Ex1-7 Layer II-VI cortical neurons ^33^, whereas Cluster n3-n5 each mapped to specific sets of interneuron subclusters (Extended data Fig. 5b). The n4 subcluster corresponded to PVALB^+^ and SST^+^ layer IV-VI (In6-8) cells. The n5 subcluster mapped to layer I/II/VI VIP^+^ interneurons (In1-3), and the n3 subcluster mapped to a mixture of NDNF^+^ and CCK^+^ interneuron subsets (In1, In4-5). Conversely, clusters n1 and n6 separated according to disease status and contained a mixture of inhibitory and excitatory neurons, mostly corresponding to Ex3 Layer IV excitatory neurons and CALB^+^, PVALB^+^ and SST^+^ In6-8 interneurons. The AD n1 subcluster was enriched for autophagy, and responses to various stimuli including hormones, lipids, topologically incorrect protein (Fig. 3e-h), exemplified by specific expression of multiple heat shock proteins (*i.e., HSPA1A, HSPB1, HSPA1B, HSPH1, HSP90AB1, HSP90AA1, HSPA8* and *HSPA5*). The control n6 subcluster was enriched in ribosomal (*RPS* and *RPL* families) and oxidative phosphorylation pathways (mitochondrial complex I-V genes), suggestive of high energy requirements and usage by these neurons ^41^. Together these data may indicate that the molecular identity of specific neuronal subsets is more susceptible to AD-related changes, specifically responses to foreign stimuli, consistent with the reported resistance of deeper layer neurons (layer V-VI) to the toxic effects of amyloid_42_.

In the case of oligodendrocytes, our data revealed that while the control oligodendrocyte subclusters (o5 and o6) are composed of a mixture of cells originating from all samples (Fig. 3i-j), the AD-specific o1-3 subclusters distinctly cluster by sample and are segregated in both UMAP space and in terms of ontological features (Fig. 3i-l). These differences may indicate that interindividual genetic variation across AD patients or slight differences in regional sampling of entorhinal cortex may affect oligodendrocyte signatures to a greater extent than observed for other cell types, where individual subclusters generally comprised cells from multiple samples.

### Cell subcluster-specific activity and regulation of GWAS candidate genes in human AD brain

Large scale genetic screening studies of AD susceptibility, such as GWAS and exome sequencing, have provided a large repertoire of pathological pathways and candidate gene targets ^4, 43^, and have highlighted the microglial genetic vulnerability in AD ^1, 2, 44^, spurring an entire field of inquiry. Here, we leveraged our single cell data resource to examine cell type specificity of expression of about 1,000 GWAS candidate genes for AD and AD-related traits (Supplementary Table 4, ^45^). To assess cell-type specificity, using the Expression Weighted Cell Type Enrichment (EWCE) package^44^, we calculated the specificity score of each GWAS gene associated with AD, LOAD, AD biomarkers, and AD neuropathologic change. Briefly, the specificity score is a measure of the proportion of a particular gene’s expression across all cell types. Therefore, a high gene specificity score for a cell type indicates a high proportion of that gene’s expression being captured in that particular cell type. First, corroborating earlier evidence, a number of previously described microglia-specific GWAS genes showed highly specific expression in microglia in our dataset, including *INPP5D, HLA−DRB5, PLCG2, HLA−DRB1, CSF3R* and *MS4A6A* (Fig. 4a, *left panel*). In addition, we detected two microglia-specific GWAS genes not previously associated with microglia, *RIN3* and *TBXAS1* (Fig. 4a, *left panel*). The former is involved in microglial endocytosis and interacts with *BIN1* and *CD2AP*, suggesting possible involvement in APP trafficking ^46, 47^. The latter is a potent vasoconstrictor in cerebral circulation, which has been linked to ischemic stroke ^48^. Notably, TBXAS1 is also the target of two FDA-approved drugs for stroke and peripheral vascular disease, dipyridamole ^49^ and picotamide ^50^, respectively, and dipyridamole was shown to inhibit amyloid-induced microglial inflammation ^51^. Therefore, our data showing significant microglia-specific *TBXAS1* expression (FDR<10^-3^) prompt further detailed functional investigations in a microglial model to study the gene’s contribution to AD, potentially in context of cardiovascular risk ^52^.

Functionally, LOAD associated GWAS genes were enriched for inflammatory, phagocytic/endocytic pathways, lipid metabolism and synaptic and axonal function ^4^. A number of neuron-specific GWAS genes have been linked to neuronal migration (e.g.*, NRG1*) and synaptic transmission/adhesion (*GABRG3, PCDH8, GRM1, RIMBP2* and *CDH9*; Fig. 4a). Interestingly, two of the neuronal-associated GWAS genes are involved in RNA splicing pathways (*SRRM4, ELAVL2*; Fig. 4a, *left panel*), as aberrant splicing was recently shown to be associated with AD in a combined aging cohort of 450 patients ^53^.

Next, we compared our single-cell level differential expression data in AD with previous bulk microarray data from the hippocampus and neocortex of AD patients ^54, 55^ (Fig. 4a, *right panels)*. *ADAMTS18*, a GWAS hit previously identified as specific to oligodendrocytes, showed cell-type independence with upregulation in AD microglia, astrocytes, oligodendrocytes, and OPCs, but downregulation in neurons. Other concordant changes of GWAS genes were dominated by a single cell type, for instance, upregulation of *MYT1* in AD oligodendrocytes, *MS4A6A* in AD microglia, *KCNN3* in AD astrocytes as well as downregulation of *VSNL1* in AD neurons and *RGS20* in AD astrocytes. Not unexpectedly, a large majority of previously identified GWAS DEGs between AD and control patient brains displayed cell-type specific expression patterns in our data, further highlighting the information value of single-cell analyses in human AD.

We further explored the functional relevance of GWAS genes using two paradigms: (*i*) we first examined subcluster-specific changes in expression of top AD GWAS genes (Fig. 4b-g and Supplementary Table 5), and (*ii*) we integrated GWAS genes into transcription factor (TF)- driven regulatory modules to understand the drivers of cell fate transitions in AD (Fig. 5). Within each cell type, the top AD GWAS genes for each subcluster were chosen based on their absolute log fold changes with respect to all other subclusters. Importantly, our single cell data allowed us to uncover sets of GWAS genes with divergent cell-type specific expression in AD cell subclusters. For instance, we observed cell-type specific regulation of *APOE* which was downregulated in AD OPC (O1), oligodendrocyte (o2) and astrocyte subclusters (a1,a2), while it was upregulated in a microglial AD subcluster (m1). Our results are in line with recent reports that reduced *APOE* expression in *APOE4* isogenic iPS-derived astrocytes resulted in impaired lysosomal functions and amyloid clearance ^56^, while increased *Apoe* expression in microglia has been associated with AD phenotypes (^25, 26^, Grubman et al., accompanying manuscript). To our knowledge, no prior study has examined the effects of *APOE* expression or genotype on oligodendrocyte or OPC function. However, given the important role of oligodendrocytes in synthesis and secretion of cholesterol ^57^, it is possible that loss of endogenous *APOE* expression in OPCs or oligodendrocyte subclusters may impair myelination, and indeed MRI analyses in 104 healthy patients showed an allele-dependent effect of *APOE* on myelin breakdown ^58^. Together, our data uncover cell subcluster-specific expression patterns of *APOE* in human AD brains, the combined and cell-specific consequences of which warrant further examination in iPSC-derived and mouse models of AD pathogenesis.

We also detected instances where GWAS genes showed coordinated up- or down-regulation in multiple AD cell subclusters. After *APOE*, the most strongly associated genetic risk factor for LOAD, *BIN1*, was specifically increased in AD subclusters a1 and n1 (Fig. 4c-d), consistent with previous reports of increased *BIN1* in AD brains ^59^. Both clusters were enriched for genes relating to autophagy and responses to incorrectly folded proteins (Fig. 3d,h), in line with the endocytic function of *BIN1* in amyloid processing ^60^ and reported direct interaction with tau^59^. However, *BIN1* has 10 transcripts, the relative abundances of which cannot be resolved from 3’ biased 10X data. Different *BIN1* isoforms exhibit specific functions in endocytosis, calcium signalling and apoptosis, and distinct expression patterns of these transcripts were associated with greater or reduced amyloid load in AD patients ^61^. *FRMD4A*, a LOAD GWAS gene ^62^ which we found to be expressed in all cell-types, showed concordant downregulation in subclusters comprised of AD patient cells in microglia, astrocytes and oligodendrocytes (Fig. 4b, c, e). FRMD4A interacts with the Arf6 and PAR complex to regulate cell polarity, but has only been studied in primary cortical neurons, where *FRMD4A* loss enhanced tau secretion ^63^.

An intriguing observation is that control subclusters also displayed higher expression levels of AD GWAS genes (*i.e., APOE* was upregulated in a4/a8, *APOC1* upregulated in m4, and *HLA- DRB1* was upregulated in m5), which may represent differential susceptibility of particular functional subpopulations to gene variants identified by GWAS (Fig. 4b-c). Together, these data highlight the advantage of studying single-cell data in order to understand the effect of disease gene variants on cell subtype-specific genetic susceptibility, and may explain why conventional (whole-body) rather than conditional/cell-type specific gene knockouts in AD models have often yielded discrepant results.

### The master lysosomal regulator, TFEB, drives a network of GWAS genes that controls cell fate transition to AD in astrocytes

To examine the causal relationships governing subcluster specific functional changes, we next systematically examined the TF dynamics underlying subcluster-specific cell state transitions towards AD. To this aim we used CellRouter ^64^ to predict TFs that drive transitions from control to AD subclusters. Briefly, CellRouter constructs path trajectories across single cell subpopulations and builds gene regulatory networks (GRNs) using mutual information, a metric drawn from information theory that measures the amount of information one random variable gives about another. Subsequently, GRN scores were calculated to reflect how well a TF and its downstream targets correlate with the identified trajectory. CellRouter did not detect any TFs for the endothelial subpopulation possibly due to the insufficient number of endothelial cells for trajectory detection. For all other cell types, our analysis unveiled common TFs predicted to control multiple transitions within a cell type, such as *AEBP1* (Fig. 5a), with previously reported upregulation in AD brains ^65^ and association with amyloid pathology ^66^. *AEBP1* displayed a high GRN score for transitions from every control astrocyte subcluster towards a1 AD astrocytes. Our analysis identified several other TFs with high GRN scores for multiple lineage conversions to AD subclusters including *SOX10*, *MYRF* and *NKX6−2*, predicted to regulate generation of n1, o1/o2, a1 subclusters (Fig. 5a), with functional enrichment of glial and neuronal development, myelination and unfolded protein response (Extended data Fig. 5c).

We set out to investigate the relationships between TFs and their regulation of GWAS genes associated with AD, highlighting relevant GRNs containing GWAS genes as downstream targets (Fig. 5b). These GRNs had the highest average log fold change across their GWAS-associated downstream targets. Integration of TF-driven regulatory modules with downstream GWAS gene targets revealed subcluster-specific control of AD cell fate transitions. For example, we discovered that *HIF3A*, a TF that inhibits hypoxia-induced gene expression ^67^, initiated a regulatory cascade containing multiple AD GWAS genes (*NPAS3, BIN1, MOBP, CLDN11* and *ADARB2*) that drives cell fate from n6 to n1. In addition, *HIF3A* was also involved in the transitions towards o1, a1-a2, m1 and O1 subclusters (Fig. 5b). However, different functional gene networks downstream of *HIF3A* were predicted for individual cell fate transitions (Extended data Fig. 5d). The vascular associations for AD and the reported effects of peripheral hypoxia on amyloid clearance from the brain ^68^, warrants further investigation of the output of cell-subcluster specific responses to hypoxia. Our analyses also showed that the *TFEB* gene, which was upregulated in diseased astrocytes^69^, acts upstream of 10 GWAS loci for AD (namely: *BIN1, CLDN11, POLN, STK32B, EDIL3, AKAP12, HECW1, WDR5, LEMD2* and *DLC1*), which are also dysregulated in AD astrocytes (Fig. 5c). Notably, this upregulated regulatory module, which was enriched in chaperone mediated responses to unfolded protein and MHC complex binding (Fig. 5d), underlies the cellular transitions from control to AD states in specific astrocyte sub-populations, which are otherwise missed in bulk tissue expression data alone. These results establish a functional link between specific astrocyte sub-populations and AD, and suggest that the contribution of multiple GWAS loci to AD susceptibility is coordinated and mediated by TFEB activity in these astrocytes.

Thus, our data and analyses provide functional annotations of several cell-type specific TF-driven regulatory networks, which will help to elucidate the cellular context and regulatory mechanisms for manifold susceptibility loci identified in AD GWAS.

### An online single cell atlas of human AD brain transcriptomes

Above we provided examples on how our data resource can be leveraged to investigate single genes and regulatory networks at single cell resolution in AD brain, and how it can be integrated with external resources (*e.g.*, GWAS data) to gain deeper insights into regulatory disease mechanisms. To facilitate AD researchers to access and mine our data resource beyond the standard GEO source repository, we provide an interactive online tool to visualise and interrogate our dataset with numerous explorative analyses: including to display the expression of any gene of interest in all individual cells, to examine cell identity, cell heterogeneity, the differential gene expression and ontologies between subclusters, and to find novel markers of cell types and subclusters.

In particular, the web interface is split into four tabs/sections. In the first tab, users can interrogate the dataset at a global level involving all the cell types in three ways. Firstly, the user can plot the gene expression using dimension reduction projections such as UMAP or PCA while simultaneously visualizing cell metadata including cell type, disease state and sequencing statistics (Fig. 6a,b). Secondly, the user can explore and investigate gene expression patterns of multiple genes in individual cell types or disease states using various visualizations (bubble plots, heatmaps and violin plots; Fig. 6c). Finally, the first tab also allows the user to tabulate the proportion of cells in different groupings (Fig. 6d). In the second tab, users can interrogate the dataset by the individual cell types in a similar fashion as the first tab. In the third tab, users can query and download the differential expression results at the level of individual cell types and subclusters, as well as disease state comparisons generated as described in the methods, and the associated gene set enrichment analyses. In the fourth tab, users can query the TFs predicted to regulate cell fate transitions between any subclusters of the same cell type. Taken together, we believe this comprehensive resource will facilitate AD research by providing a platform to generate a plethora of hypotheses related to AD cell identity, diversity and pathogenic mechanisms and to identify novel cell-specific pathways and targets.

**Fig. 6.**
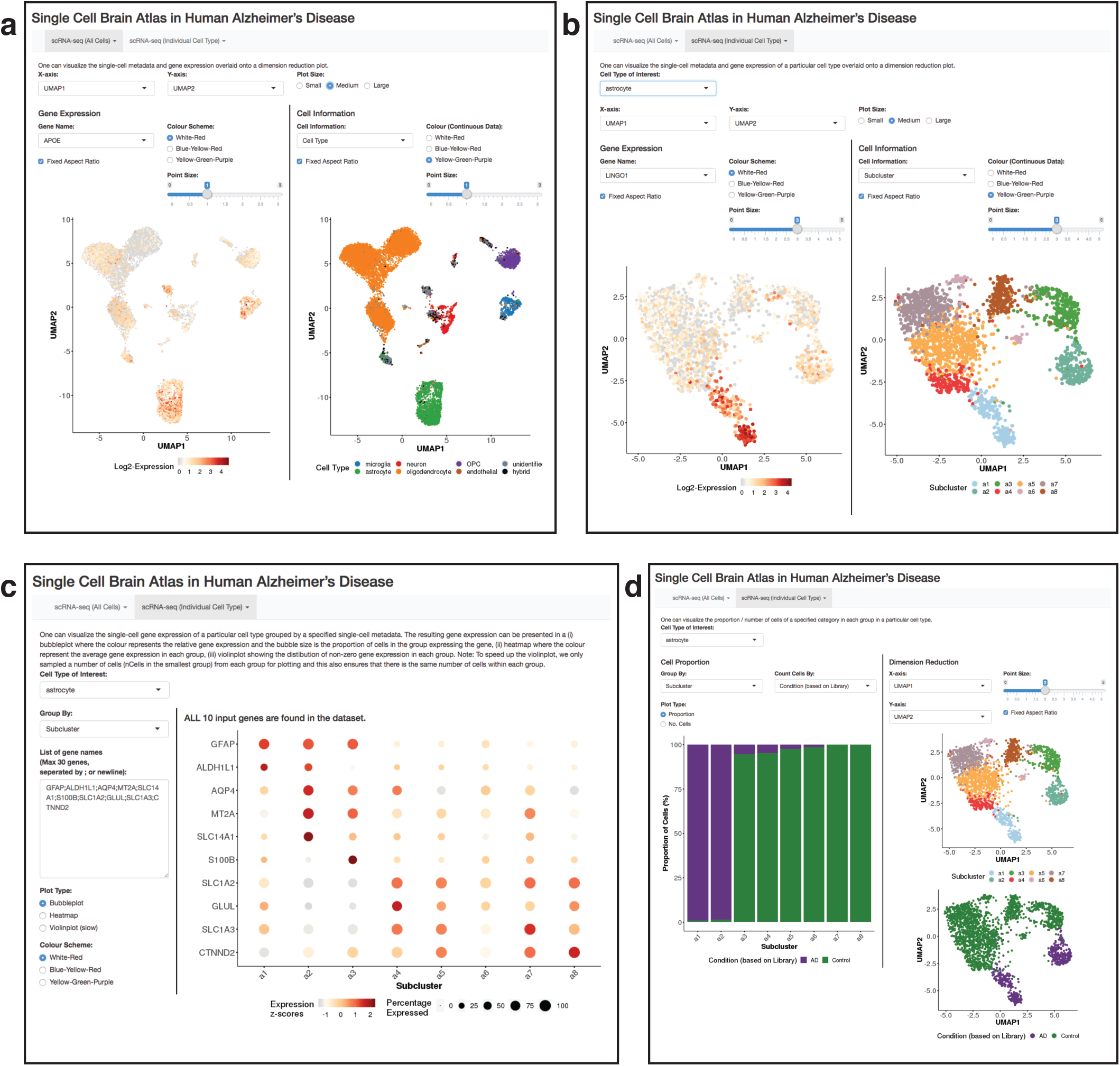
A user-friendly web resource of single nuclei transcriptome data analysis in human AD. Screenshots from the web resource including **a**, left, dimensionality reduction visualisation of expression of any gene of interest in all individual cells, and right, visualisation of single cell metadata by UMAP or PCA, including cell type, # UMI per cells, # genes detected per cell, % mitochondrial genes, AD or control cells, library ID, and cell specificity scores. **b**, subcluster dimensionality reduction plots, showing astrocyte subclusters as an example, left, showing single cell gene expression, and right, subcluster composition. **c**, single-cell gene expression of a particular cell type grouped by specified single-cell metadata, that can be viewed as a bubble plot, violin plot or heatmap. In the bubble plot, the colour represents the relative gene expression and the bubble size is the proportion of cells in the group expressing the gene. **d**, Cell proportion plot of subclusters visualized by library in astrocyte subclusters.

## Discussion

To the best of our knowledge, our data represent the first single cell transcriptomes of AD patient brains, providing a unique resource for future studies seeking to understand cellular heterogeneity and define functional changes at single-cell resolution in human AD brain. To demonstrate the utility of this resource in aiding to uncover the genetic mechanism underlying cellular heterogeneity and function in AD, we have provided several examples of applications and data analyses, including integration with external datasets related to genetic susceptibility to AD. By doing this, we have been able to gain several novel insights into cell subtype specific regulation and cell identity changes in AD.

In the accompanying paper by Grubman et al. 2019, we identified a microglial XO4^+^ signature linked to plaque phagocytosis and regulated by *HIF1A* that increases the phagocytic potential of microglia in a mouse model of AD. Here, using our human single cell sequencing resource we demonstrated that the XO4^+^ signature is molecularly conserved in human AD microglia (specifically to a subpopulation of microglia, m1, see Fig. 5a, Extended data Fig. 4a-d, accompanying paper Fig. 5a-h). This provides a concrete example of cross-species conservation of cell-type specific molecular signatures associated with functional phenotypes in AD brain. Similarly, we believe our resource will facilitate investigators to translate the findings generated in various model systems and experimental models of AD to human AD.

The identification of transcription factors that orchestrate the conversion of control to AD cell signatures in the human brain can pinpoint specific molecular processes and mechanisms for therapeutic intervention. Accounting for the intrinsic cellular complexity in AD brain, we provide a detailed map of all TF-driven conversions from control to AD across all cell subpopulations in the human brain (Fig. 5). The systematic annotation of these regulatory networks in AD brain will prompt experiments and testable hypotheses to elucidate disease mechanisms at the single cell level. Given the cellular complexity observed in AD pathophysiology, knowledge of the specific cell population(s) where a given regulatory program is operating to drive disease will help the design of more robust and effective (i.e., cell-type specific) cellular studies and therapeutic approaches.

The many disease-associated genes identified by GWAS remain largely orphan in regards to a specific functional context in human AD brain. Our data resource allowed us to provide direct insights into the cells and ontological cell subclusters in which these disease genes operate in human AD brain. Notably, our data reveal complex patterns of expression changes for multifold AD genes within or across specific cell populations. This complexity should be taken into account to enhance our interpretation of genetic discoveries in AD. For example, our data on cell-type specific expression of GWAS genes will prompt expression quantitative trait loci (eQTL, ^1^), splice QTL (sQTL, ^53^) and single-cell ATAC-Seq analyses ^70^ tailored to specific cell populations, to inform functional specialization of AD gene variants and “regional” (*i.e.*, cell-type specific) susceptibility to disease.

Finally, our resource allowed us to explore the genetic contributors to AD beyond analysis of single gene effects in specific cell types and/or subpopulations. Specifically, integrated gene networks and GWAS data at the single-cell level to detail the coordinated (TF-driven) contribution of multiple GWAS genes to AD susceptibility in specific cellular transitions from control to AD states. For example, we show how the TFEB transcription factor, which is upregulated in diseased astrocytes, acts upstream of 10 GWAS loci for AD and which are also dysregulated in astrocytes from AD patients’ brains. Functionally, this TFEB-network of GWAS genes was enriched for chaperone mediated responses to unfolded protein and MHC complex binding, and our data show how this process underlies the cellular transitions from control to AD states in specific astrocyte sub-populations. These results provide insights into disease networks in the human brain, and uncover molecular interactors that relate to driving nodes (*e.g.*, TF) connecting pathways to genetic susceptibility (*e.g.*, GWAS genes) to disease in specific cell populations.

We anticipate that our resource will stimulate and allow further discoveries in many different areas (Table 1) as exemplified by our work.

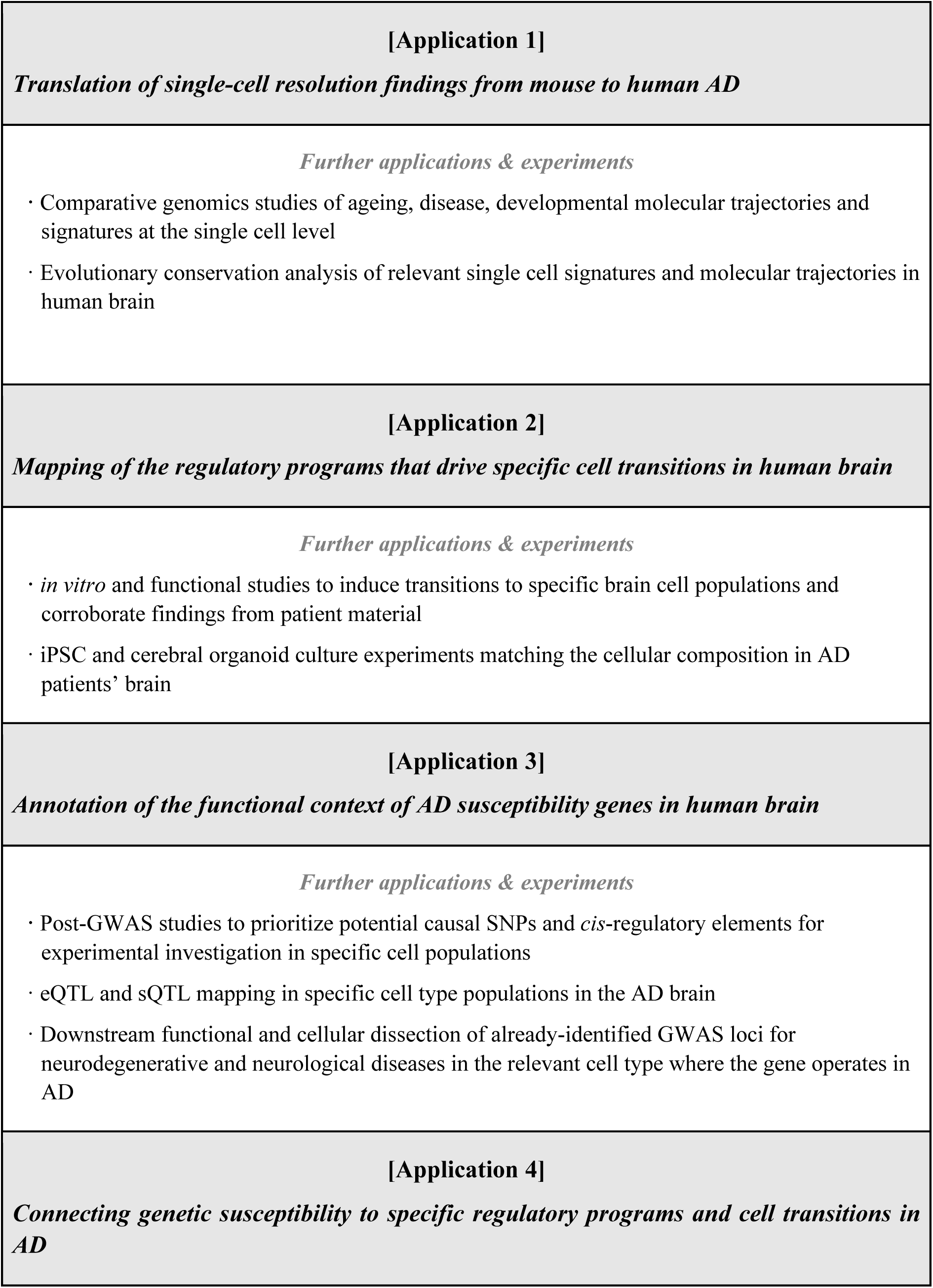
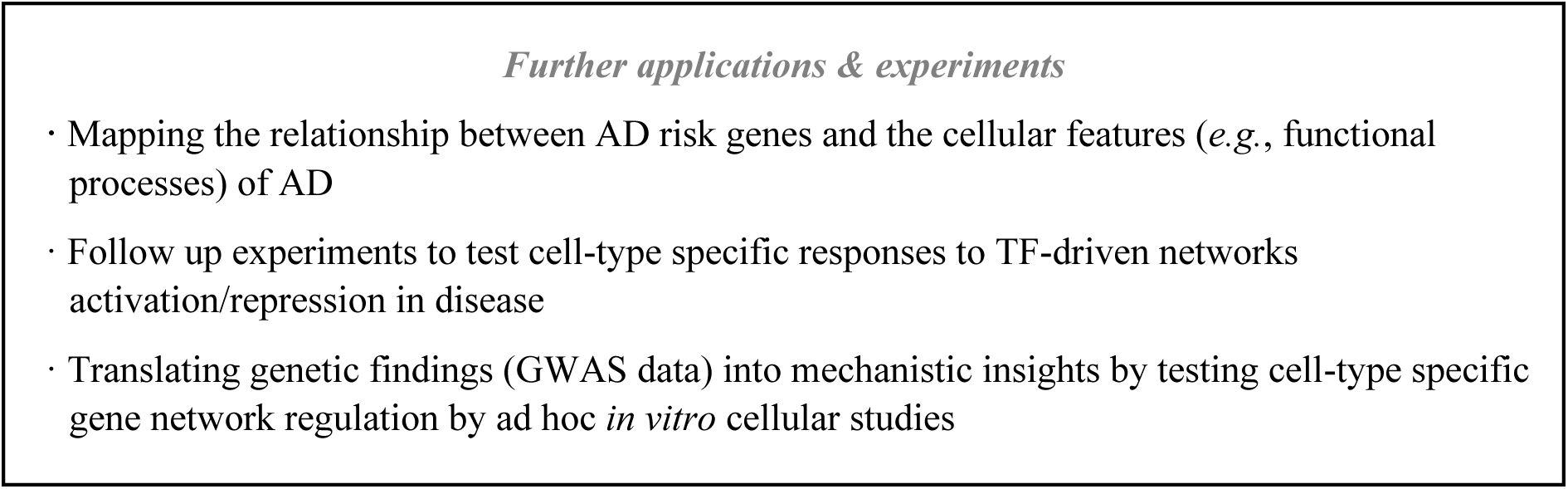

## Methods

### Nuclei isolation sorting from human AD brain tissue

Entorhinal cortex tissue from post-mortem Alzheimer’s disease and non-disease aged matched individuals was obtained from the Victorian Brain Bank (Ethics Approval: MUHREC 2016-0554; patient demographics in Supplementary Table 1). Nuclei isolation was carried out using the Nuclei Isolation Kit: Nuclei EZ Prep (Sigma, #NUC101) as described in ^9^. Briefly, tissue samples were homogenized using a glass dounce grinder in 2 ml of ice-cold EZ PREP and incubated on ice for 5 min. Centrifuged nuclei (500 × g, 5 min and 4 °C) were washed in ice-cold EZ PREP buffer, and Nuclei Suspension Buffer (NSB; consisting of 1× PBS, 1% (w/v) BSA and 0.2 U/μl RNase inhibitor (Clontech, #2313A). Isolated nuclei were resuspended in NSB to 10^6^ nuclei per 400 μl), filtered through a 40 μm cell strainer and counted with Trypan blue. Nuclei enriched in Nuclei Suspension Buffer were stained with DAPI (1:1000) for nuclei isolation using the Influx cell sorter (BD Biosciences, Franklin Lakes, NJ; 70 μm nozzle, 21-22 psi). Nuclei were defined as DAPI^+^ singlets. Sorted nuclei were counted twice prior to loading onto the 10X Chromium (10X Genomics). Library construction was performed using the Chromium Single Cell 3′ Library & Gel Bead Kit v2 (10X Genomics, #PN-120237) with 18 cDNA pre-amplification cycles and sequencing on one high-output lane of the NextSeq 500 (Illumina).

### Mapping single nuclei reads to the genome

Using the Grch38 (1.2.0) reference from 10x Genomics, we made a pre-mrna reference according to the steps detailed by 10x Genomics (https://support.10xgenomics.com/single-cell-gene- expression/software/pipelines/latest/advanced/references). Cellranger *count* was used to obtain raw counts. Our single nuclei data initially comprised 8 10x runs consisting of 4 AD runs and 4 control runs. Each run had 2 patients (see patient information above). 2 runs (1 AD run and 1 control run) were discarded because of high neuronal enrichment (see Extended data Fig. 2) possibly indicating neuronal contamination or technical artefacts. In all, this resulted in a final dataset of 6 10x runs consisting of 3 AD runs and 3 control runs.

### Quality Control for expression matrix

The raw expression matrix was composed of 33,694 genes and 14,876 cells. Genes without any counts in any cells were filtered out. A gene was defined as detected if 2 or more transcripts were present in at least 10 cells. 100 PMI-associated genes, as defined by Zhu et al in the cerebral cortex, that were detected in our data were removed ^71^.

For cell filtering, cells outside the 5th and 95th percentile with respect to number of genes detected and number of unique molecular identifiers (UMI) were discarded. In addition, cells with more than 10% of their UMIs assigned to mitochondrial genes were filtered out. The matrix was normalized with scale factor of 10000 as recommended by the *Seurat* pipeline ^34^ before *FindVariableGenes* was used to define variable genes with the parameters x.low.cutoff = 0.0125, x.high.cutoff = 3, and y.cutoff = 0.5. *ScaleData* was used to center the gene expression. Overall, the resulting filtered matrix consisted of 10,850 genes and 13,214 cells.

### Cell Type Identification

BRETIGEA ^12^ is a R package which utilizes brain cell-type marker gene sets curated from independent human and mouse single cell RNA datasets for cell type proportion estimation in bulk RNA datasets. It can also be employed in single cell RNA datasets for the identification of cell types. For the reference datasets, BRETIGEA uses well-annotated and well-referenced datasets from ^16^ and ^17^. The 6 main cell type lineages identified by BRETIGEA are neurons, astrocytes, oligodendrocytes, microglia, oligodendrocyte progenitor cells (OPCs), and endothelial cells. We obtained the gene sets from BRETIGEA for all 6 cell types and calculated a module score for each cell type using Seurat’s *AddModuleScore* function. This function calculates the average expression levels of each cell-type gene set subtracted by the aggregated expression of a background gene set. Cell type identification was then performed in 2 steps.

Firstly, each cell was assigned a cell type based on the highest cell type score across all 6 cell types. Furthermore, we defined a cell as a hybrid cell if the difference between the first and second highest cell type scores are within 20% of the highest cell type score:

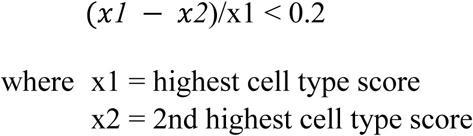

Secondly, for each cell type, we assumed normality of the gene score distribution and applied z-score transformation. Subsequently, in order to consider poorly identified cells i.e. cells with low cell type scores, for each cell type, cells with low cell type score (5th percentile and below) were relabelled as “unidentified” cells.

Alternatively, these cells may represent cells in which a true cell signature is more difficult to determine due to a large number of gene dropouts: indeed, a large proportion of hybrid and unidentified cells are from the AD15042_04076 library corresponding to the 2 oldest AD patients, where we detected only a median of 473 genes per cell.

Overall, we obtained 449 microglia, 2,171 astrocytes, 656 neurons, 7,432 oligodendrocytes, 1,078 OPCs, 98 endothelial cells, 925 unidentified cells, and 405 hybrid cells. From Uniform manifold approximation and projection (UMAP) visualization, the clusters separate well based on cell type, supporting the accuracy of our cell type identification method.

### Human single nuclei UMAP and clustering analysis

Seurat was used for normalization, scaling, and finding variable genes in the same manner as described in *“Quality control for expression matrix”*. For each cell type, Seurat’s *SubsetData* was used to subset the data. Subsequently, each cell type undergoes the same procedure of normalization, scaling, and finding of variable genes. Prior to UMAP calculation, Principal Component Analysis (PCA) was first performed to obtain a small number of principal components (PCs) as input to the UMAP algorithm. The number of PCs to be included varies across the individual cell types and the global dataset and was determined using the Elbow method. The percentage variance explained by each PC was plotted and the number of PCs was then chosen at the “elbow” of the plot where a substantial drop is observed in the proportion of variance explained. Seurat’s *FindClusters* was used to derive subclusters within each cell type with resolution 0.8. The number of principal components for each cell type used are below:

| Cell Type | Number of Principal Components |
| --- | --- |
| All Cell Types | 19 |
| Microglia | 15 |
| Astrocytes | 16 |
| Neuron | 15 |
| Oligodendrocytes | 15 |
| OPC | 16 |
| Endothelial | 18 |
| Unidentified | 7 |
| Hybrid | 9 |

### Human single nuclei differential expression and gene set enrichment analysis

Differential expression was performed using the empirical Bayes quasi-likelihood F-tests (QLF) in the edgeR package (v3.20.8). Genes are deemed significantly differentially expressed if they have FDR < 0.05. Several differential expression comparisons were performed. First, to determine cell type specific genes, differential expression was performed between the cell type of interest and the average of the remaining five other cell types. For example, to compare the first cell type with the average of the remaining cell types, the argument *contrast=c(1,-0.2,-0.2,-0.2,- 0.2,-0.2)* was supplied to the *glmQLFTest* function in edgeR. Second, to identify transcriptomic differences between AD patients and control patients at the cell type level, differential expression was performed between cells from AD patient libraries and control patient libraries for each cell type. Third, we designated subclusters to be either AD/control/undetermined based on the cell composition. A subcluster is designated to be AD is more than 80% of the cells originate from an AD patient and the same applies for assigning control subclusters. Otherwise, a subcluster is designated to be undetermined. Differential expression was then performed between cells in the AD subclusters and control subclusters for each cell type. Fourth, to identify subcluster specific genes within each cell type, differential expression was performed between the subcluster of interest and the average of the remaining subclusters of the same cell type. Fifth, differential expression was also performed between pairs of subclusters of the same cell type. For all the above-mentioned differential expression, gene set enrichment analysis was also performed using the fgsea package (v1.4.1) with 100,000 permutations.

### Obtaining novel cell type markers

We first obtained DE genes between each cell type and all 7 other cell types. For each cell type, we overlapped DE genes with FDR < 0.05 and log FC > 2 with the human marker genes obtained from BRETIGEA. The non-intersected genes are deemed the novel cell type markers.

## Cell Router Analyses

### Gene Regulatory Network Analysis

CellRouter is a single-cell RNA sequencing algorithm used to identify gene regulatory changes along transitions between user-defined cell states ^64^. CellRouter analysis was run using the pipeline provided by the original authors (https://github.com/edroaldo/CellRouter) as of 14th December 2018. For each cell type, we input raw counts into CellRouter. This was followed by CellRouter’s built-in normalization, scaling, PCA calculation, and t-SNE (t-distributed stochastic neighbor embedding plot) visualization^72^. t-SNE was performed with perplexity of 20 and 1000 iterations. For each cell type, a KNN (k - Nearest Neighbor) network was built with CellRouter’s *buildKNN* using jaccard similarity with parameters k = 10 and number of principal components as derived from the table in *Single Human Nuclei Analyses.* CellRouter’s *findPaths* was used to find the trajectory from one subcluster to another. Paths were processed with CellRouter’s *processTrajectories* by minimising path cost and setting minimum number of cells in trajectory and neighbors to 2 and 3 respectively. To identify genes regulated along each trajectory, CellRouter’s *correlationPseudotime* was performed using spearman correlation and using only the top 75th (positively correlated) and bottom 25th percentile(negatively correlated) to define our gene networks in the next step. CellRouter’s *smoothDynamics* and *clusterGenesPseudotime* were subsequently performed. Gene regulatory networks (GRNs) were constructed using *buildGRN* with z-score threshold of 1.65. With respect to the GRN scores calculated by CellRouter, only the top 90th and bottom 10th percentile gene regulatory networks were taken as the final outputs from the CellRouter pipeline. “Hs” was used as the species input. Igraph R package was used within CellRouter’s pipeline for processing networks (http://igraph.org).

### Pruning of Gene Regulatory Networks (GRNs)

Importantly, for each GRN, we noticed that the downstream targets identified by CellRouter might not be differentially expressed (DE). Therefore, we pruned each network by removing non-DE downstream targets. Next, for each transition, we selected GRNs containing the DEGs that are most highly DE genes. We first calculated a gene score for each downstream target by multiplying absolute log2(Fold Change) and −log10(false discovery rate). For each transition, we removed a GRN from our analysis if its gene scores were not significantly higher than the gene scores of all the other DE genes for that respective transition via a one-sided Wilcoxon non-parametric test. This ensured that the remaining GRNs have downstream targets with gene scores significantly higher than the gene scores of the DE genes which are not part of each respective GRN. In other words, for each trajectory, the remaining GRNs have downstream targets representing the most highly DE genes. Lastly, we ensured that the transcription factors identified were indeed human transcription factors by overlapping them with the list from Lambert et al ^73^. It is important to note that after these post-processing and pruning steps, some transitions did not have any significant TFs, perhaps suggesting that the transition is between subclusters which have similar transcriptional landscapes.

### Visualization of GRN Scores and Networks

For figure 4a, pheatmap ^74^ was used to plot up to the top 5 GRN scores for each trajectory. For this visualization, we only considered TFs which are inducers i.e. upregulation of TFs leads to upregulation of downstream targets. ggplot2 ^75^ was used for visualizing the GRN scores for each trajectory in the R shiny App. Cytoscape ^76^ was used to construct the network, and yFiles was used to construct the hierarchical network layout. Cell symbols were created with BioRender (https://app.biorender.com). For each cell type, networks plotted are the networks with the highest average logFC of GWAS hits taken from the differential expression between the source subcluster and each of its target subcluster. These GWAS hits are colored by the average log2 FC.

## Single-cell level annotation of GWAS candidate genes

### Calculating GWAS Specificity Scores

GWAS data were downloaded from the NHGRI-EBI catalog ^45^ on the 25th of February, 2019. Specifically, data for four traits were downloaded - “Alzheimer’s Disease” (EFO-0000249), “Alzheimer’s Disease Biomarker Measurements” (EFO-0006514), “Late-Onset Alzheimer’s Disease” (EFO-1001870), and “Alzheimer’s Disease Neuropathologic Change” (EFO-0006801); see Supplementary Table 4 for list of genes. We first removed gene duplicates and GWAS loci in intergenic regions, and used a p- value ⩽ 9×10^-6^ to identify significant associations. Then, since GWAS signals can point to multiple candidate genes within the same associated locus, for each AD-related phenotype we choose to focus on the “Reported Gene(s)” (i.e., genes reported as associated by the authors of each GWAS study). Overall, we retrieved 980 GWAS-reported genes for downstream analysis. For calculation of cell-type specificity scores, we followed the Expression Weighted Cell Type Enrichment (EWCE) method described by Skene et al ^44^. Briefly, this approach involves the following steps. First, ANOVA was performed to remove uninformative genes whose expression do not vary significantly across cell types. We implemented a p-value threshold of 0.00001 for the F-statistic, which is the default value used in the EWCE R package. Second, specificity scores were calculated using *generate.celltype.data*. Finally, we defined a GWAS-associated gene as cell-type specific if its specificity score is at least at the top 99th percentile of the specificity score distribution across all cells with respect to all the GWAS-associated genes. Using this criterion, we identified 30 cell-type specific genes, whose average expression (LogCPM) across cell types is in Fig. 4a.

## Comparison of single cell data to bulk AD gene expression profiles

### Comparing with public datasets

Blalock et al ^55^ performed microarray analyses for laser-captured hippocampal tissue from control and AD patients and generated a list of genes and corresponding directionality of expression in AD. “UP” genes were given a logFC of +1 and “DOWN” genes were given a logFC of −1. We overlapped this list with the 30 cell-type specific GWAS genes provided that these genes are differentially expressed between subcluster-driven AD and control clusters (FDR < 0.10). For both the microarray and our datasets, we normalize the logFC by the respective maximum absolute logFC before visualization. We also compared our data with Tan et al ^54^ who performed microarray analysis in the neocortex of AD patients. We overlap the DE genes identified by Tan et al ^54^ (FDR < 0.10) with our 30 cell-type specific GWAS genes. Once again, genes are considered differentially expressed between subcluster-driven AD and control clusters if FDR is less than 0.10. LogFC were normalized using the same methodology described earlier in this section.

### Finding Subcluster specific GWAS Hits

After obtaining the differentially expressed genes (FDR < 0.05) between each subcluster and the other subclusters for each cell type, we overlapped these genes with the 980 GWAS-reported genes (Supplementary Table 4). For visualization in Fig. 4b, up to the top 3 genes with the highest absolute logFC for each subcluster were plotted for each cell type.

## Functional annotation of GRN

### Functional annotation of TFEB networks

Enrichment of the pruned TFEB networks for specified transitions - a3 to a1, a6 to a1, a7 to a1, and a8 to a1 - was performed using *moduleGO* from the R packages DGCA ^77^ and GOstats ^78^. The top 5 Gene Ontology (GO) terms for each transition were plotted in Fig. 5d. The conserved TFEB network represents the genes that are common across all 4 transitions. Reported p-values are adjusted for multiple testing by Benjamini-Hochberg (BH) method ^79^.

### Functional annotation of top GRN networks

Enrichment of the pruned GRN networks was performed using the same procedure described in “*Functional annotation of TFEB networks*”. The top 5 GO terms for each GRN network were plotted in extended Fig. 5c.

## Comparison with Lake et al neuronal data

Processed neuronal expression data from ^33^ were downloaded from https://hemberg-lab.github.io/scRNA.seq.datasets/human/brain/. Briefly, Lake et al utilized single nucleus sequencing on post-mortem human brains from 6 different regions and identified 16 neuronal subtypes. The original log CPM expression matrix consists of 25,051 genes and 3,042 cells. For each of the 16 neuronal subtypes, we calculated the average log CPM for each gene to construct a 25,051 by 16 reference expression matrix. We projected our neuronal data onto this reference expression matrix using methods as described in Reference Component Analysis (RCA) ^80^. In short, RCA pipeline involves the calculation of the pearson correlation coefficient to find out how closely associated the projected data is to the reference data. The visualized outputs are the z-score transformations of the correlation coefficient raised to the fourth power.

## Comparison with Lin et al and Liddelow et al astrocyte data

Astrocyte data were downloaded from ^35^. Briefly, Lin et al. identified 5 astrocyte subpopulations in human adult brain. Each of these 5 reported astrocyte subclusters has its own module or set of marker genes (FDR < 0.10). For each module, we converted mice gene names to human orthologs using *biomaRt (2.34.2)*. In addition, for each module, we removed genes which appear in other modules, thereby giving us specific genes for each module. We plotted the average expression of each module in our 8 astrocytic subpopulations. For visualization, we normalized the expression of each module by the maximum module expression across all 8 astrocytic subpopulations. Similarly, we plotted the average expression of the pan-reactive, A1-specific, and A2-specific gene modules as identified in ^24^ across all 8 astrocytic subpopulations.

## Shiny App development

*R shiny web application:* To allow for easy visualization of the extensive dataset and analysis involved in this work, a web interface (http://adsn.ddnetbio.com) was created using the shiny package (v1.1.0).

## Data availability Statement

All single-cell RNA-seq sequencing data are available from the Gene Expression Omnibus (GEO) upon the acceptance of the manuscript. Data can be visualized via the interactive web application at adsn.ddnetbio.com.

## Code availability Statement

Code is available from the authors by reasonable request.

## Competing Financial Interests

The authors declare no competing financial interests.

## Acknowledgements

The authors acknowledge Flowcore and Micromon, Monash University, for the provision of instrumentation, training and technical support. The Australian Regenerative Medicine Institute is supported by grants from the State Government of Victoria and the Australian Government. A.G was funded by a NHMRC-ARC Dementia Fellowship and Dementia Australia Research Foundation Grant. J.M.P. was funded by a Sylvia-Charles Viertel Fellowship. Part of this work was funded by a Monash Network of Excellence grant. G.S. was funded by the Yulgilbar Foundation.

## Author Contributions

AG and JMP conceived the study and designed experiments and together with EP and OJLR designed the bioinformatics analysis. AG, GS and XYC performed nuclei isolation and FACS. CM performed pathological assessment of human control and AD cases. GC and EP performed GWAS integration, CellRouter and network analyses. JFO and OJLR performed cell type and cell subcluster identification, DGE and GSEA analyses. JFO and OJLR developed the shiny web interface. JP, RS, SB, DVL and RL worked up the protocol for single nuclei sequencing from human brain. AG, GC, JFO, OJLR, EP and JMP wrote the manuscript. All authors approved of and contributed to the final version of the manuscript.

## Supplementary Tables

**Supplementary Table 1 Patient pathology and clinical history.**

**Supplementary Table 2 Novel cell type marker genes.**

**Supplementary Table 3 Genes relating to AD KEGG Gene set enrichment in individual subclusters.**

**Supplementary Table 4 Genes associated with AD and AD-related traits by GWAS, grouped by GWAS trait (AD, LOAD, biomarkers, neuropathologic change)**

**Supplementary Table 5 DEGs overlapping with GWAS genes**

## Notes

http://adsn.ddnetbio.com

