## Extended data Fig for "A single cell brain atlas in human Alzheimer’s disease"

### Extended data Fig. 1

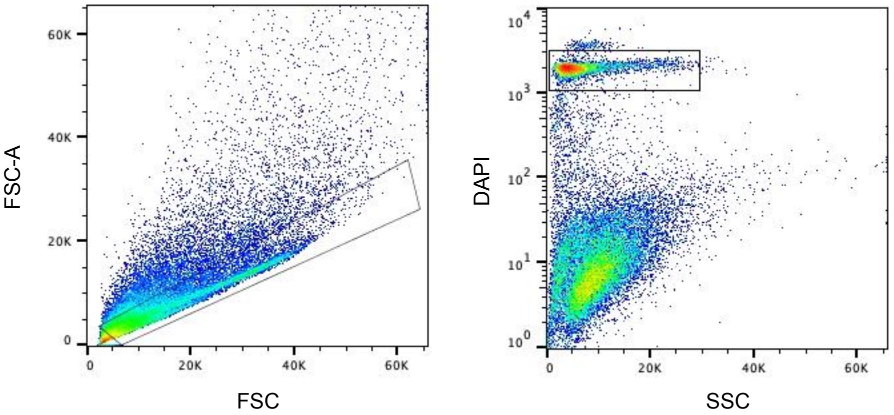

**Extended data Fig. 1 Gating strategy for FACS isolation of single nuclei.** Nuclei isolated using the EZ-Prep kit from entorhinal cortex of AD and control patients were FACS sorted on BD Influx using 70 µm nozzle, 21-22 psi. Single $DAPI^+$ events were considered nuclei.

### Extended data Fig. 2

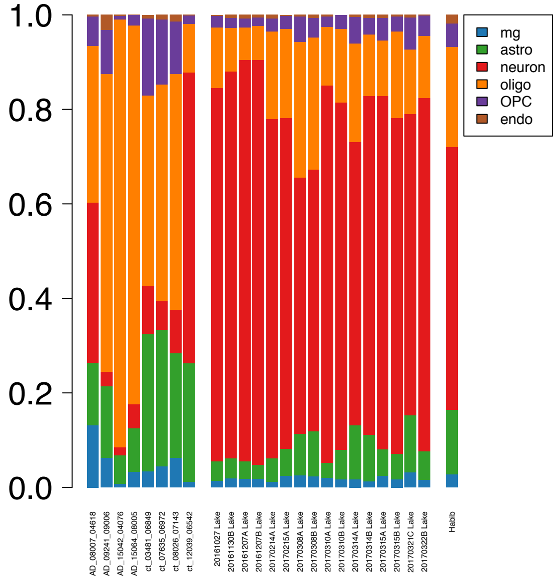

**Extended data Fig. 2 Single nuclei metadata. a,** cell proportion plots by library for 8 libraries showing exclusion of 2 libraries due to high neuronal count and comparison of cell type proportions recovered in this study compared to cell proportions in other single nuclei studies [33] and [9]. Cell proportions are shown by sequencing library, where this information is available. **b,** library proportion plots for cell subclusters

### Extended data Fig. 3

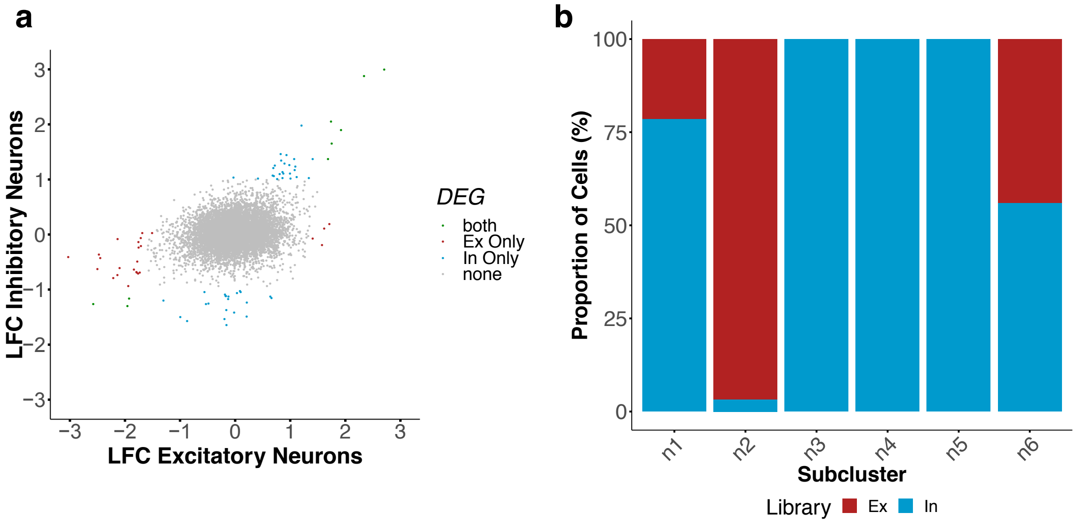

**Extended data Fig. 3 AD related changes in excitatory and inhibitory neurons.** Scatter plot showing the relationship between cell-specific and common DEGs in control and AD, excitatory and inhibitory neurons. Cutoff for significance is |LFC|=0.5

### Extended data Fig. 4

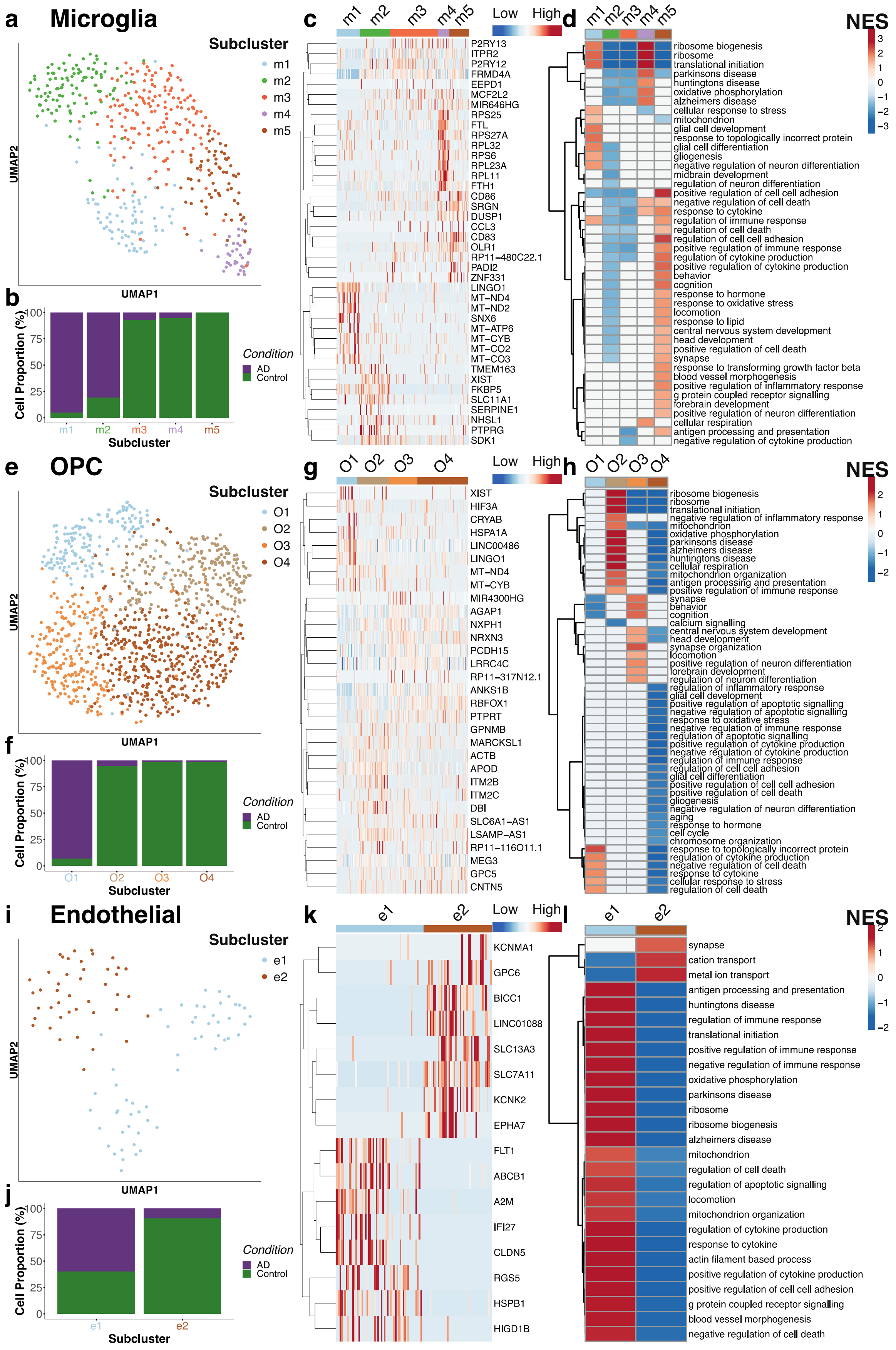

**Extended data Fig. 4 Single nuclei sequencing of human AD and control entorhinal cortex reveals homeostatic, AD-specific and shared ontological cell subclusters. a,e,i,** UMAP visualization of subclusters of microglia (**a**), OPCs (**e**) and endothelial cells (**i**) showing **b, f, j**, the composition of cells in subclusters by disease state, **c, g, k,** hierarchical clustering and heat map colored by single cell gene expression of subcluster-specific genes (top 8 genes were shown per cluster). **d, h, l,** GSEA of subcluster specific genes coloured by normalised gene set enrichment scores for the gene ontologies shown in each cell subcluster.

**Extended data Fig. 5**

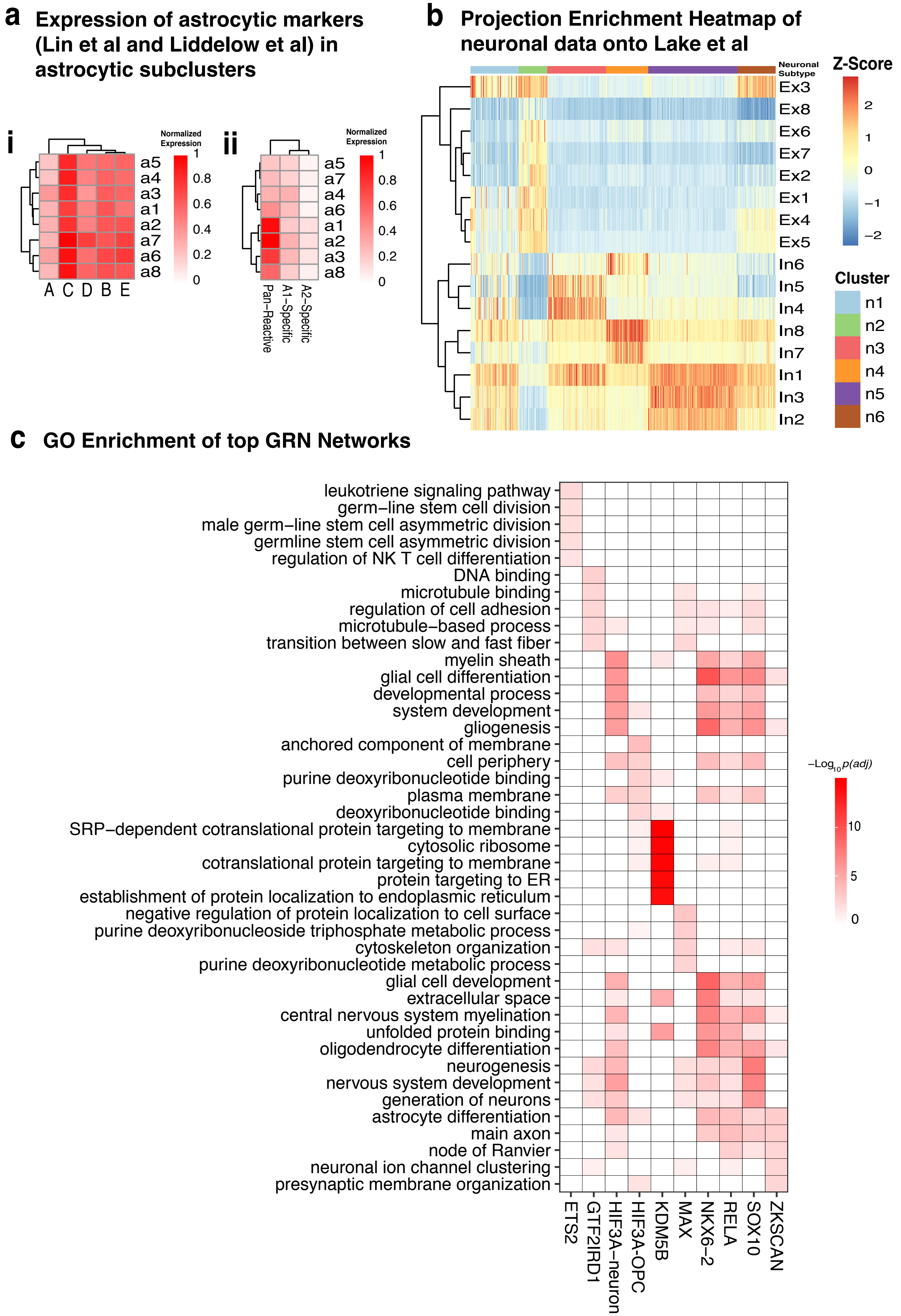

**Extended data Fig. 5** Comparison of astrocyte signatures to $\mathbf{a_i}$[24]; $\mathbf{a_{ii}}$ [35]. **b,** Comparison of neuronal subcluster signatures to [33](Ex1-8, In1-8). **c,** Functional annotation of GRNs from Fig. 5b

**Supplementary Tables**

**Supplementary Table 1 Patient pathology and clinical history.**

**Supplementary Table 2 Novel cell type marker genes.**

**Supplementary Table 3 Genes relating to AD KEGG Gene set enrichment in individual subclusters.**

**Supplementary Table 4 Genes associated with AD and AD-related traits by GWAS, grouped by GWAS trait (AD, LOAD, biomarkers, neuropathologic change)**

**Supplementary Table 5 DEGs overlapping with GWAS genes**
